# Enhanced perceptual task performance without deprivation in mice using medial forebrain bundle stimulation

**DOI:** 10.1101/2022.03.29.486230

**Authors:** Antonin Verdier, Noémi Dominique, Déborah Groussard, Anna Aldanondo, Brice Bathellier, Sophie Bagur

## Abstract

Training animals on perceptual decision-making tasks is essential to many fields of neuroscience. Current protocols generally use water as a reward in water-deprived animals. This can be challenging since it requires balancing animals’ deprivation level with their well-being. Moreover, trial number is limited once they reach satiation. Here, we present electrical stimulation of the medial forebrain bundle (MFB) as an alternative reward in mice that avoids deprivation entirely while yielding stable motivation for thousands of trials. We trained MFB rewarded mice to perform a series of auditory discrimination tasks using either licking or lever pressing as a behavioural report. MFB animals learnt tasks at similar speed to water-deprived mice and more reliably reached higher overall accuracy in harder tasks. Moreover, they performed up to 3800 trials within a session without loss of motivation. Importantly, MFB stimulation does not impact the sensory behaviour under study since psychometric parameters and response times are preserved. Finally, in accordance with the lack of deprivation, MFB mice lack signs of metabolic or behavioural stress compared to water-deprived mice (weight loss, open field, home cage behaviour). Overall, using MFB stimulation as a reward is a highly promising tool for task learning since it enhances task performance whilst avoiding deprivation.

## Introduction

Probing the sensory abilities and decision-making processes of animals in order to model these faculties and link them to neural activity is central to systems neuroscience. This relies on training animals to perform a range of instrumental tasks. In mice, the most common protocols water deprive animals to motivate them to lick for small water rewards. Although this approach has enabled multiple successful studies, it suffers from certain experimental drawbacks. First, water deprivation requires careful and time-consuming monitoring of water consumption in order to avoid discomfort to animals while maintaining their motivation. Despite well-established procedures, motivation levels can be hard to control depending on the individual animal, the housing conditions or the experimental environment, leading to variable results. Finally, once animals reach satiation they stop performing, thus limiting the number of trials per session. This can impede the possibility of probing fine psychophysics [1] and curtail analysis through lack of data, particularly the ability to link noisy neural data to behaviour and perception.

Here we propose a novel reinforcement method which addresses these issues by using intracranial stimulation of the Medial Forebrain Bundle (MFB) as a reward. After the remarkable observation that stimulation in the hypothalamic region caused rats to return to the location where they had received the stimulation [2], a flurry of research identified the MFB as the region evoking maximal self-stimulation without adverse effects [3]. This fibre bundle courses via the lateral hypothalamus between multiple regions of the brain’s reward system, most notably the ventral tegmental area and the nucleus accumbens. This self-stimulating effect has mainly been used in order to study the neural underpinnings of motivation itself but some studies have used this rewarding effect to perform spatial conditioning [4]–[6]. This suggested to us that this method could provide a reward system without the aforementioned drawbacks of satiation or prior deprivation.

Here we demonstrate that MFB stimulation can be used in head-fixed mice to flexibly train animals on auditory discrimination tasks with high and stable performance over thousands of trials per session. We compare performance and learning with that of mice trained on the same tasks using water deprivation and find evidence for similar learning rates as well as more reliable learning on hard tasks. Moreover, we show that MFB stimulation reduces physiological and behavioural stress in comparison to water deprivation. Therefore, training with MFB stimulation can contribute to reducing the impact of experiments on animal welfare, when compatible with the underlying scientific question. Overall, this novel technique provides an easy-to-implement alternative to water deprivation with demonstrable experimental advantages.

## Results

### MFB rewarded mice can learn auditory discrimination tasks

The MFB is a long fibre tract of around 400μm in diameter which can be targeted with standard stereotaxic procedures for stimulation with a bipolar electrode (Fig. S1). The implantation surgery is short (∼1 hour) (Fig. 1A) and the small size of the electrode allows to easily combine MFB stimulation with head-posts, large cranial windows (Fig. 1B) and/or electrophysiology. In some mice (n=16), we tested the rewarding effect of MFB stimulation using a conditioned object preference assay (Fig. S2G-I, Video 1) in which stimulation was manually elicited when mice contacted a cotton swab in the cage. All mice showing conditioned preference for the cotton swab were classified as having a “functional MFB”, defined as successful acquisition of the report behaviour for the perceptual task (defined in Materials and Method, Fig. S2). This first step allows an efficient triage of animals after surgery. Post-hoc histology showed that absence of conditioning was due to large electrode targeting errors (>400μm, Fig. S1). However, small targeting errors (<400μm) had little impact on the rewarding effect of the stimulation and on success of the MFB-driven discrimination task (Fig. S1). This was an important factor for the high success rate of our implantation surgeries. Amongst all implanted mice in our lab, 87% (82/94) experienced functional MFB-based reinforcement (Fig. S2 A,D). Detailed breakdown of mice included in all experimental groups is given in Table S1.

**Figure 1.**
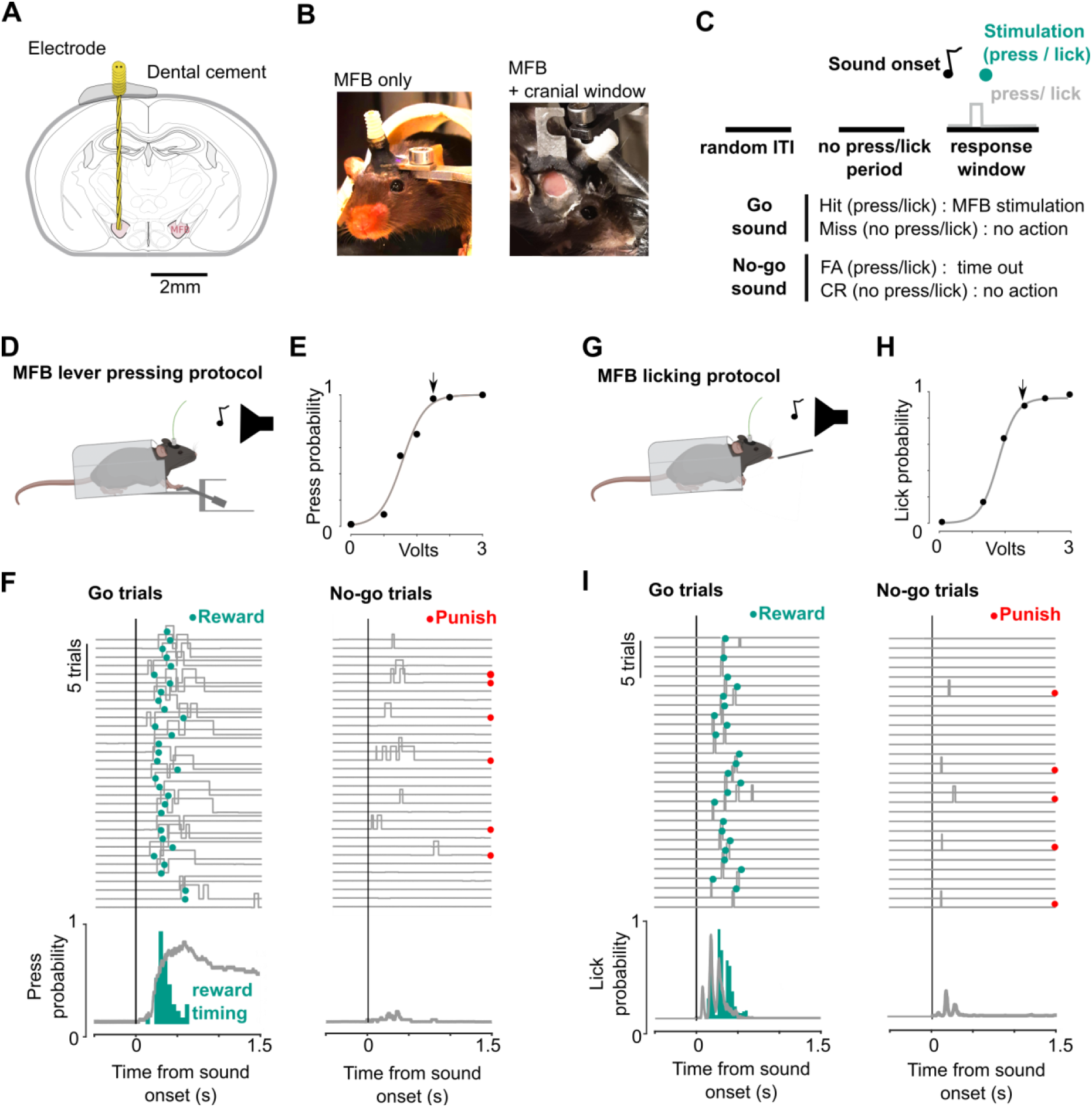
Protocol for perceptual discrimination learning with MFB reward. A. Medial forebrain bundle implantation site of bipolar electrode fixed with dental cement to skull B. Photo of head-fixed mouse showing head post and MFB electrode (left) and mouse with 5mm cranial window over the auditory cortex, head-post and MFB electrode (right) illustrating that the small MFB electrode can be combined with recording methods. C. Structure of the Go-No-go step of the discrimination task D. Scheme of head fixed mouse during task : pressing the lever with the forepaw after Go sound elicits stimulation via the implanted electrode E. Calibration curve for an example mouse. Measured probability of pressing the lever during 1.5s response window after Go sound for MFB stimulation of different voltages (blue dots) and sigmoid curve fit to data (grey line). Arrow indicates voltage used for learning. F. Traces of lever press for 30 Go and 30 No-go trials from an example mouse after training (top) and the mean lever press trace for all trials (bottom). For each Go trial, the time of reward (50ms after lever press) is shown by a green dot and for each No-go trial, trials punished by a time out are indicated by a red dot. The distribution of reward timings is shown in green. G.H.I As in D.E.F for licking

In this study, we established that MFB stimulation can be used as a reinforcement signal for sound discrimination learning (Fig. 1C) using two types of reporting motor actions often used in sensory discrimination tasks: the pressing of a lever or the licking of a spout (Fig. 1D-I). The first step of the training (Free reward) was to elicit MFB stimulation as soon as mice spontaneously licked or pressed the lever in order to associate report behaviour to reward (Fig. S2B,E). The second training step (Go training), was to condition reward on licking / pressing during a 1.5s window following sound presentation to associate the stimulus and the reward (Fig. S2C,F). Licking report behaviour was acquired on the first day whereas lever pressing required several sessions since it is a complex motor behaviour [7] (Fig. S2A-F). It is important to note that mice spontaneously licked for MFB reward without any prior water-deprivation to motivate licking. Voltage was initially set at 3V. However to optimise learning, once basic behavioural report was acquired, we calibrated the minimal voltage that elicited robust response, and used this calibrated level during subsequent training (Fig. 1E, H).

Mice were then trained on the Go/No-go step of the task (Fig. 1F,I). They had both to respond to the Go sound and to refrain from pressing following a No-go sound. False alarms were punished by a 5 to 7s timeout. Each trial was separated by a random interval during which mice had to restrain from pressing to initiate the next trial. (See videos 2 and 3 for mice performing licking or lever pressing report). These steps, along with the detail of the auditory tasks discussed below are shown in Fig. S3. In order to facilitate the use of this protocol, we supply on our Github page (https://github.com/AntoninVerdier/MFBasic) designs for the 3D models as well as instructions and code to implement a simple version of the task.

### Fast and reliable learning with MFB reward

In order to assess learning using the novel MFB procedure, we trained mice on three auditory discrimination tasks (full protocol presented in Fig. S3). To provide a reference for comparison, we also trained mice on the same tasks, using a classical water deprivation protocol. Water-deprived mice were trained to lick for a reward. MFB-deprived mice were trained either to lick or to press a lever to get the reward. Stimulus contingencies, training steps, experimenters and training apparatus were otherwise identical. In all tasks the valence of the sounds was counterbalanced across mice.

We first trained the three groups on a classical auditory frequency discrimination task (PTvsPT). Mice were trained to discriminate between two pure tones of low and high frequency (6 kHz and 16 kHz respectively). The task was considered to be learnt when accuracy was above 80% over a full session. No difference in the speed of learning was observed between the three groups, either in terms of the number of trials (Fig. 2A, Fig. S4) or the number of sessions (Fig. 2D).

**Figure 2.**
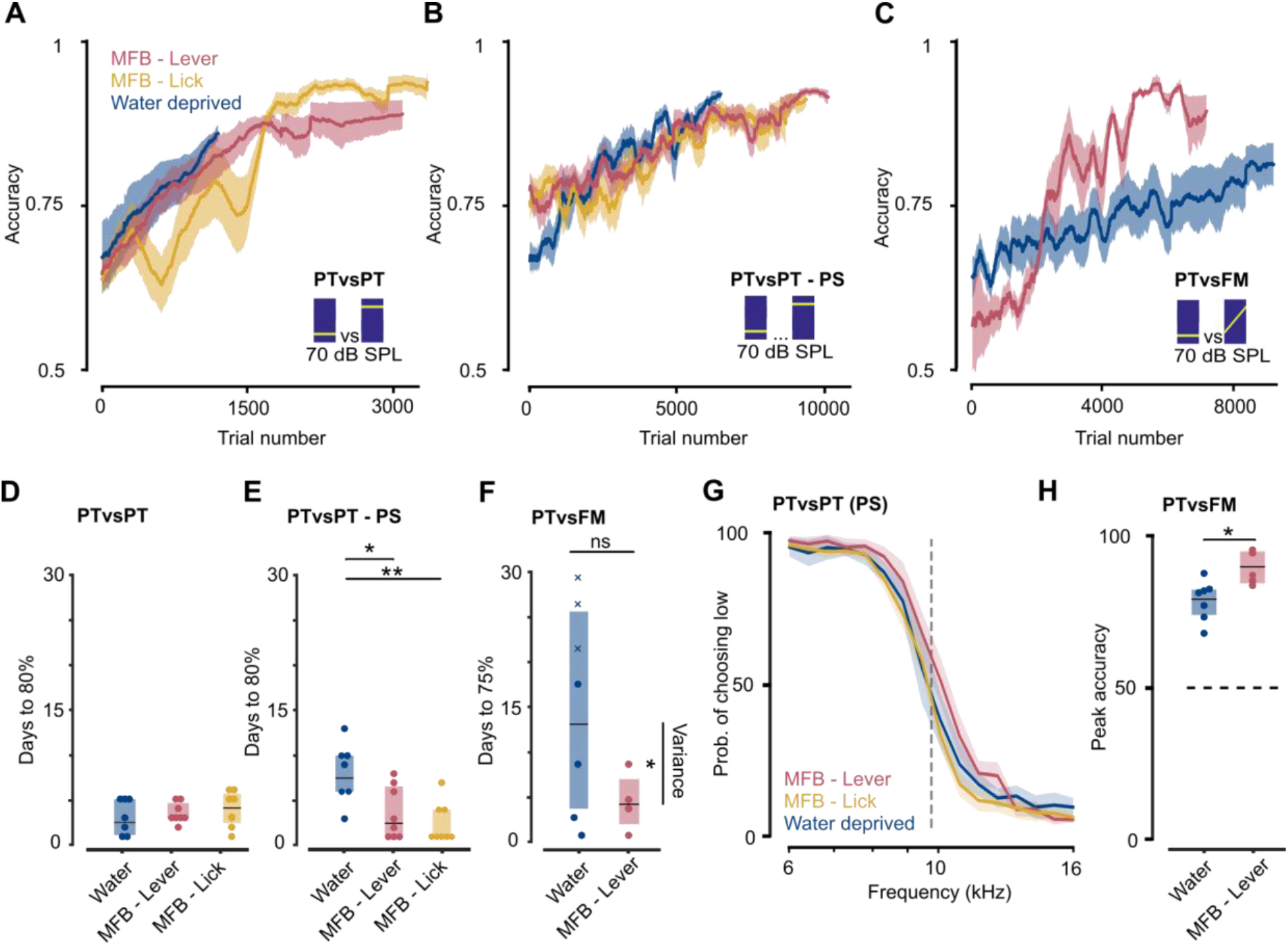
MFB-rewarded mice learn tasks at equal rates to water-deprived mice and reach higher peak accuracy on hard tasks. A.B.C. Learning curves for water-deprived (blue) and MFB-rewarded (red and yellow) mice performing the pure tone (PTvsPT) discrimination task (A -n = 7,8,8), the pure tone psychometric task (PTvsPT -PS) requiring classification of high and low pure tones (B - n = 7,8,8) and the pure tone vs frequency modulated (PTvsFM) discrimination task (C - n = 7,4). See FigS4 for independent evolution of Go vs No-go accuracy. D.E.F. Number of days necessary to reach criterion accuracy for water-deprived and MFB-rewarded mice for the three tasks as in A,B,C (Wilcoxon rank sum test, water vs MFB lever - pval : 0.74, 0.019, 0.29, ranksum statistic : 52.5, 79, 48; water vs MFB lick - pval : 0.37, 0.004, ranksum statistic: 48,75; n=7,8,8;7,8,9;7,4). The vertical bar in F indicates that variability was higher for PTvsFM task for water- deprived mice (Levene test, pval: 0.0156, Levene statistic: 8.8). For water-deprived mice that never reached criterion in the PTvsFM task, the maximum day of training is shown as a cross. G. Average psychometric function for pure tone frequency classification, drawn from the best 800 consecutive trials of each mouse for the water-deprived and MFB-rewarded mice (n=7,8,8). Shaded areas indicate SEM. H. Accuracy of the best session for water-deprived and MFB-rewarded mice for the PTvsFM discrimination task. (Wilcoxon rank sum test, pval : 0.02, ranksum statistic: 30, n=7&4)

Following learning on this first task, mice were introduced to a psychometric task (PTvsPT - PS) in which they were required to categorise 16 logarithmically-scaled sounds into low and high frequency categories (threshold frequency : 9800 Hz). As in the previous task, the learning rate did not differ between groups in the number of trials to reach the criterion. However, the number of experimental sessions necessary was higher for the water-deprived group (Fig. 2B,E) given that MFB-rewarded mice perform more trials per session (see below). All groups reached 90% accuracy and yielded the same psychometric curves (Fig. 2G). MFB therefore did not alter the measured acoustic abilities of mice that can be probed independently of the reward type once the task has been learnt.

Finally, we checked that the MFB reward could also support learning in a harder task. We trained a sub-group of 4 of the MFB lever press mice and the group of 7 water deprived mice to discriminate between a pure tone and a frequency sweep with identical start frequency (PTvsFM). This task is known to be more demanding [8] since mice cannot make a decision based on the salient sound onset but must integrate the sound over time in order to reach a decision. All MFB-rewarded mice learnt the task within 8 training sessions whereas water-deprived mice showed significantly more variable learning rates with some animals never reaching learning criterion even after 30 training sessions (Fig. 2C,F, Fig. S4). Moreover, peak accuracy was significantly higher in MFB-rewarded mice (Fig. 2H). Despite the small sample size and the difference in report behaviour (lever vs licking -see discussion), this suggests that MFB stimulation may provide more stable motivation in increasingly complex tasks, enabling faster acquisition at the cohort level.

These results indicate that the MFB stimulation technique allows for reliable learning in a range of auditory tasks with different behavioural reports.

### MFB yields stable performance for thousands of trials per session

When training relies on water rewards, satiation limits the number of trials performed in a single day or behavioural session. In our experiments, we observed a systematic decline of Go response accuracy during the session, which is likely due to satiation (Fig. 3A,B). We therefore generally limited animals to 320 trials per day and never went above 640 trials. More generally, current reports indicate that the number of trials performed in a single session by water deprived mice compatible with sustained motivation can vary between a few hundreds of trials and a maximum of 1000 trials in some cases depending on the laboratory and on protocol parameters (see literature search in Table 1).

**Figure 3.**
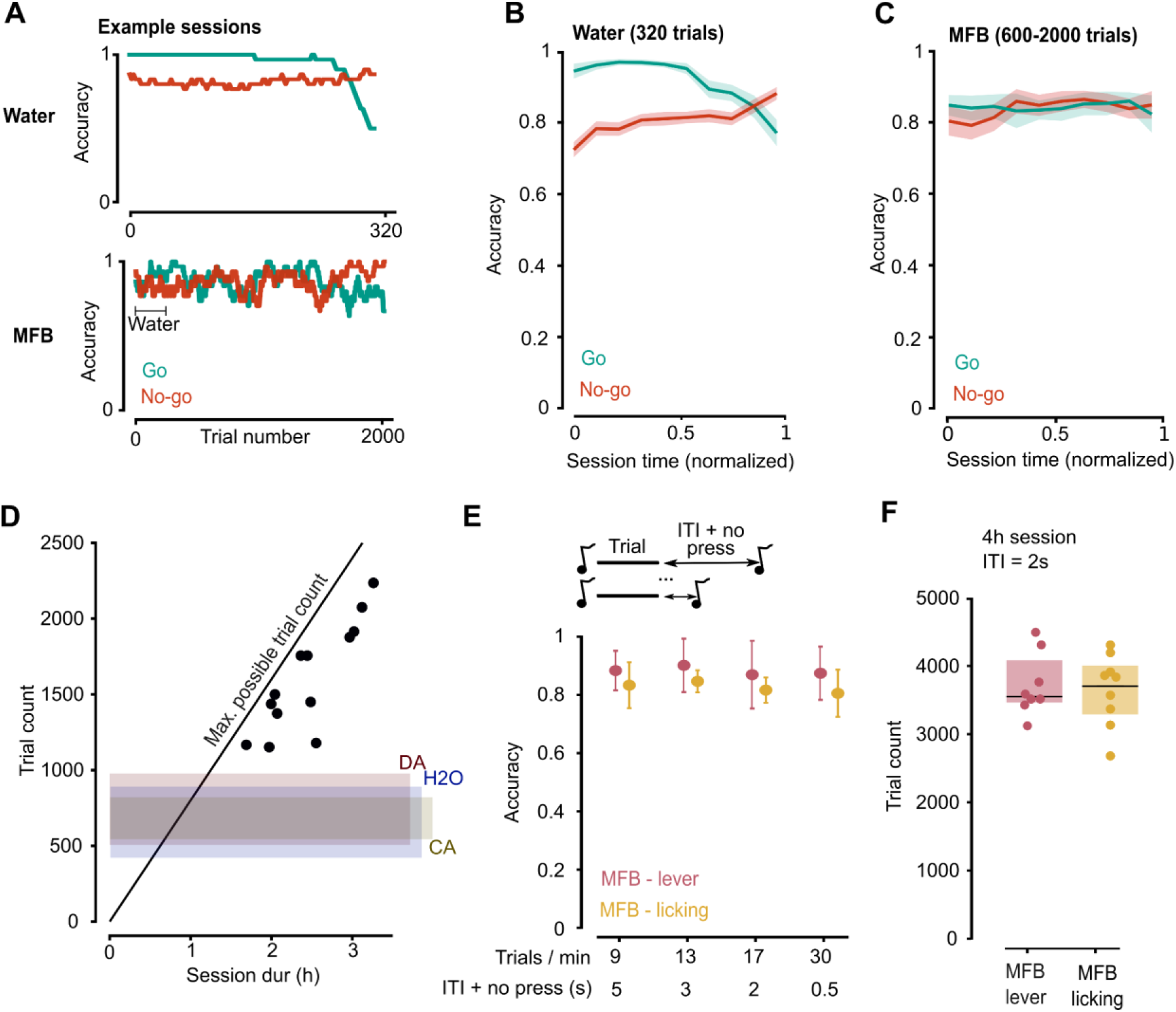
MFB-rewarded mice perform thousands of trials per session. A. Example sessions accuracy on Go (green) and No-go (red) trials for a water-deprived and a MFB leve pressing mouse in the PTvsPT task). Go accuracy drops for the water-deprived mouse after 240 trials whereas the MFB-rewarded mouse is still performing above the criterion after 2000 thousands trials. B.C. Average of Go (green) and No-go (red) accuracies for water-deprived (B) and MFB lever pressing (C) mice in the PTvsPT task. For MFB mice, in order to cumulate sessions of varying duration (600-2000 trials), time was normalised between 0 and 1 (n=7 & 8). D. Trial count of several sessions in the PTvsPT-PS task (>80% accuracy) with varying trial duration. The black line corresponds to the maximum possible number of trials mice could reach given the session duration (supposing no punishment intervals or delayed trials by licking outside of trials). Shaded areas provide the range of trial numbers found with other strategies in the literature: optogenetic stimulation of dopaminergic fibres (DA) [9] water deprivation [10], water deprivation by adding citric acid to drinking water [11]. E. Accuracy over 200 trials for different trial speeds for MFB rewarded mice performing the PTvsPT task by lever pressing or licking. Note that 500ms inter-trial intervals provide the same accuracy as 5000ms-spaced trials (n=4 & 6). F. Trial count for 4h session with 2s ITI for MFB rewarded mice performing the PTvsPT task by lever pressing or licking. Each point is one session (lever : n=8 sessions from n=6 mice, lick : n=9 sessions from n=6 mice).

**Table 1.**
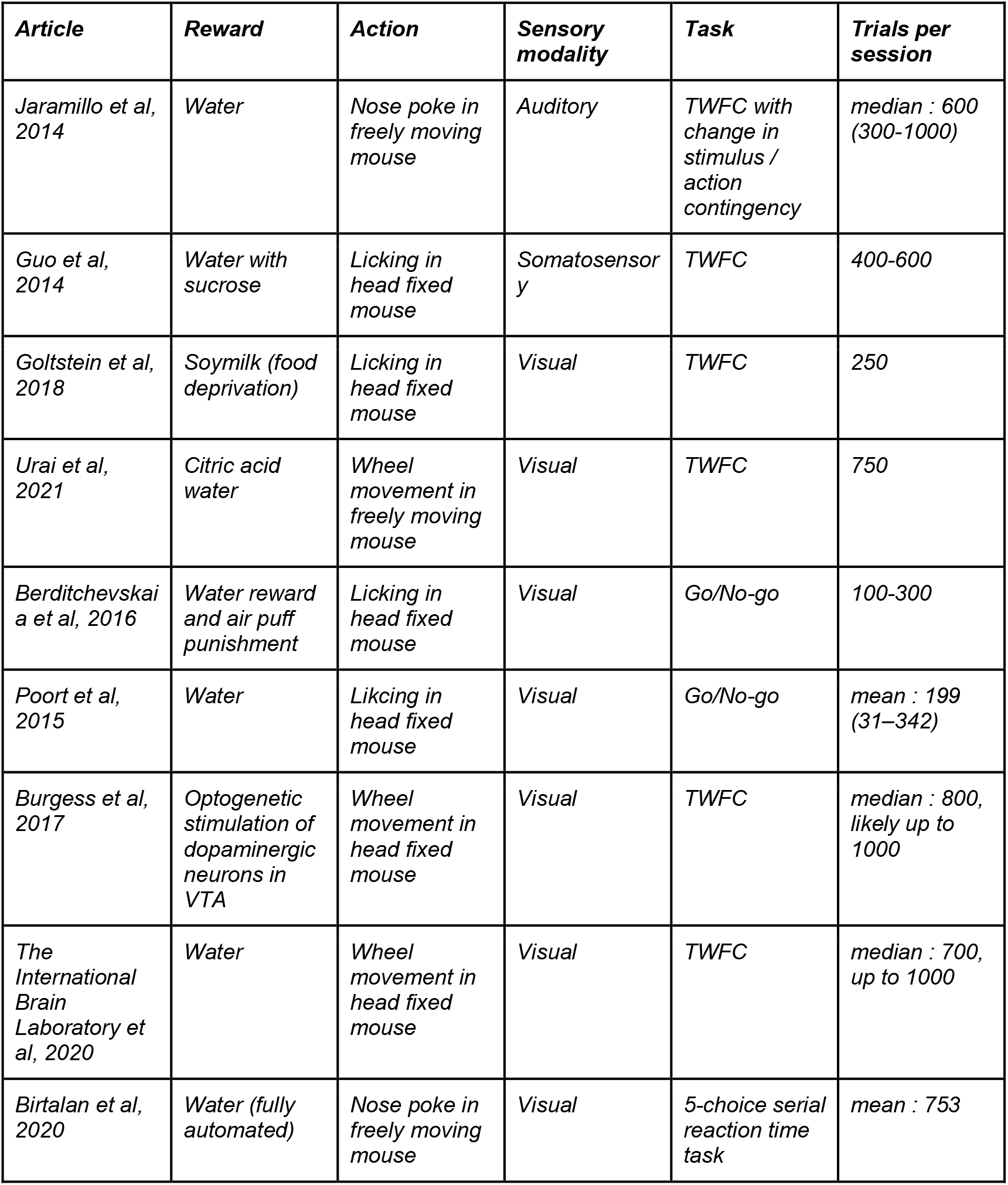
Comparison of reward systems for training in the mouse with number of trials per session

| **Article** | **Reward** | **Action** | **Sensory modality** | **Task** | **Trials per session** |
| --- | --- | --- | --- | --- | --- |
| *Jaramillo et al, 2014* | Water | Nose poke in freely moving mouse | Auditory | TWFC with change in stimulus / action contingency | median : 600 (300-1000) |
| *Guo et al, 2014* | Water with sucrose | Licking in head fixed mouse | Somatosensory | TWFC | 400-600 |
| *Goltstein et al, 2018* | Soy milk (food deprivation) | Licking in head fixed mouse | Visual | TWFC | 250 |
| *Urai et al, 2021* | Citric acid water | Wheel movement in freely moving mouse | Visual | TWFC | 750 |
| *Berdichevskaya et al, 2016* | Water reward and air puff punishment | Licking in head fixed mouse | Visual | Go/No-go | 100-300 |
| *Poort et al, 2015* | Water | Licking in head fixed mouse | Visual | Go/No-go | mean : 199 (31–342) |
| *Burgess et al, 2017* | Optogenetic stimulation of dopaminergic neurons in VTA | Wheel movement in head fixed mouse | Visual | TWFC | median : 800, likely up to 1000 |
| *The International Brain Laboratory et al, 2020* | Water | Wheel movement in head fixed mouse | Visual | TWFC | median : 700, up to 1000 |
| *Birtalan et al, 2020* | Water (fully automated) | Nose poke in freely moving mouse | Visual | 5-choice serial reaction time task | mean : 753 |

Using MFB reward, we observed during training that mice did not show signs of an irreversible drop in motivation (Fig. 3A,C-D). The number of trials was not limited by the animal’s motivation but by experimental time constraints. For example, in the psychometric task (PTvsPT -PS) and for sessions ranging between 1h30 and 3h after learning (accuracy >80%), the number of trials scaled with the session duration, without loss of accuracy and reached up to 2500 trials. The number of trials per session can be further increased by reducing the inter-trial interval. Varying this parameter in trained MFB-rewarded mice during the PTvsPT task, we observed that mice can maintain high accuracy even when performing up to 30 trials/minute, with an average trial window of 1500ms and average total inter-trial interval (randomised no press window + ITI durations, see Methods) of 500ms (Fig. 3E). This is likely enabled by the high and stable motivation of the mice but also the lack of time required for reward consumption (i.e. licking and drinking of water). Using these optimised conditions, we evaluated the maximum number of trials that could be obtained in a 4h duration using a fast 2s ITI to provide a benchmark for the novel MFB technique. We found that mice performed on average 3800 trials, with some mice achieving more than 4500 trials, a far larger number than typically reported in the literature (Fig. 3D, Table 1). Mice maintained high and consistent accuracy throughout these sessions (91%+/-3% lever press, 93%+/-6% licking) (Fig. 3F, example sessions Fig. S5).

These results show that MFB allows to train mice to perform very large numbers of trials with high performance. This provides opportunities to accelerate psychometric measurements by probing, within a single session, many repeats of multiple stimulus properties or eventually to decrease the number of training days by increasing the number of trials per day.

### MFB provides strong motivation without excess movement, impulsivity or signs of stress

The repeated use of intracranial stimulation could lead to detrimental effects on behaviour. To address this possibility, we first assessed whether the MFB led to agitated behaviour during the task, which could result from excessive activation of the dopaminergic system. We quantified the level of body movement with a piezoelectric sensor placed under the animal as a proxy for agitation due to motor response and to reward. We contrasted hit and false alarm trials during which the mouse executed the same motor action to measure reward-induced movement. In all cases, the response led to an increase in movement that however subsided more quickly during the trial in MFB trained mice, likely because they did not initiate long lick trains for water consumption (Fig. 4A). Both water and MFB reward led to an increase in movement due to reward. This increase was however much smaller than that induced simply by the animal’s response itself and of comparable amplitude for MFB and water-rewarded mice (Fig. 4A).

**Figure 4.**
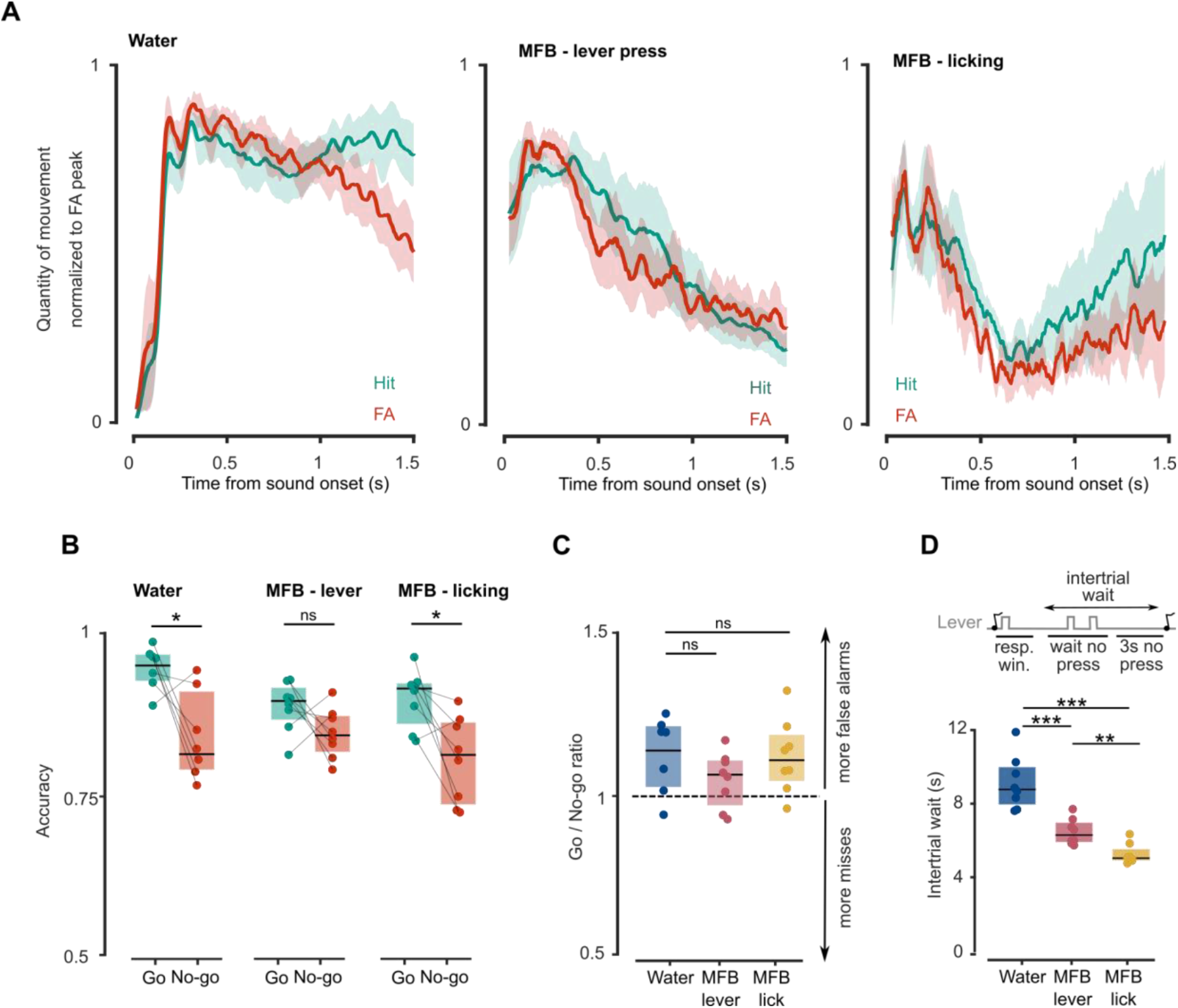
MFB stimulated mice show no signs of increased agitation or impulsivity. A Movement level for hit and false alarm (FA) trials during the psychometrics task. The difference between the two curves is indicative of the movement induced by the reward since hit trials contain movement induced by the response (licking / lever-press) and reward (water delivery or MFB stimulation) whereas FA trials only contain movement induced by the response. The movement level for each mouse is normalised by the peak of FA movement and therefore the absolute levels of movement between MFB and water cannot be compared (see materials and methods for details, n=4,8,8, error bars are sem) B. Average accuracy on Go and No-go trials in psychometrics task during the three best sessions for water-deprived (left) and MFB-rewarded lever-press and licking mice. Note that all groups are biassed towards go-responding although the effect is weaker in lever pressing MFB mice (Wilcoxon signed rank test, pval = 0.04, 0.19, 0.02 signed rank statistic : 26, 28, 34; n=7,8,8) C. Ratio of Go to No-Go accuracy showing that all groups have similar levels of impulsivity based on this measure. (Wilcoxon rank sum test, pval : 0.12, 0.77, 0.19 ranksum statistic : 70, 59, 55, n=7,8,8) D. Average wait period between trials for water-deprived and MFB-rewarded mice that captures the tendency of mice to lick / lever press during the wait period. MFB-rewarded mice are significantly more restrained during the wait period (Wilcoxon rank sum test, pval : 6.2E-4, 1.5E-4, 0.003 ranksum statistic : 98, 100, 41 n=7,8,8)

To evaluate whether MFB reward was associated with more impulsive behaviour, we measured two classical indicators of impulsivity in the Go/No-go paradigm [12]: false alarm probability, and premature responses during the inter-trial interval. As expected, mice showed worse accuracy on No-go trials than on Go trials, although this difference was not significant in MFB lever pressing animals (Fig. 4B, C). No significant difference between groups was observed for Go / No-Go accuracy ratios (Fig. 4C). However, irrespective of whether the report was licking or lever pressing, inter-trial wait periods were shorter in MFB-rewarded mice than in the water deprivation task (Fig. 4D). This indicates a reduction of pre-trial impulsivity because in our protocol the inter-trial wait period is prolonged when impulsive licks or lever press are detected (see Methods).

We also investigated signs of physiological or behavioural stress. Mice in the water-deprivation protocol underwent an initial and sustained weight loss as well as a progressive loss of weight every week before the weekend when they had one day ad-libitum access to water (Fig. 5A,B). MFB-rewarded mice however showed a trend towards increasing weight as expected by ageing mice and stable weight during the week (Fig. 5A,B). We also assessed activity levels in the homecage during the night using a running wheel. Running duration was similar in MFB-rewarded mice and in control animals left in their home cages without any training. Water-deprived mice spent less time active, as previously shown (Fig. 5C, D) [13]. To directly measure stress levels, we performed an open field test prior to any behavioural training and 12 weeks later. In both groups, we found a reduction in exploratory behaviour after training. However, reductions both in distance travelled and entries in the field centre were significantly stronger in water-deprived mice compared to MFB-rewarded mice (Fig. 5E,F). The behavioural change common to both groups may be due to the ageing of mice [14], the test-rest effect [15] or the repeated schedule of head-fixation and training although the literature suggests that head-fixation itself does not modify behaviour in the open-field [16].

**Figure 5.**
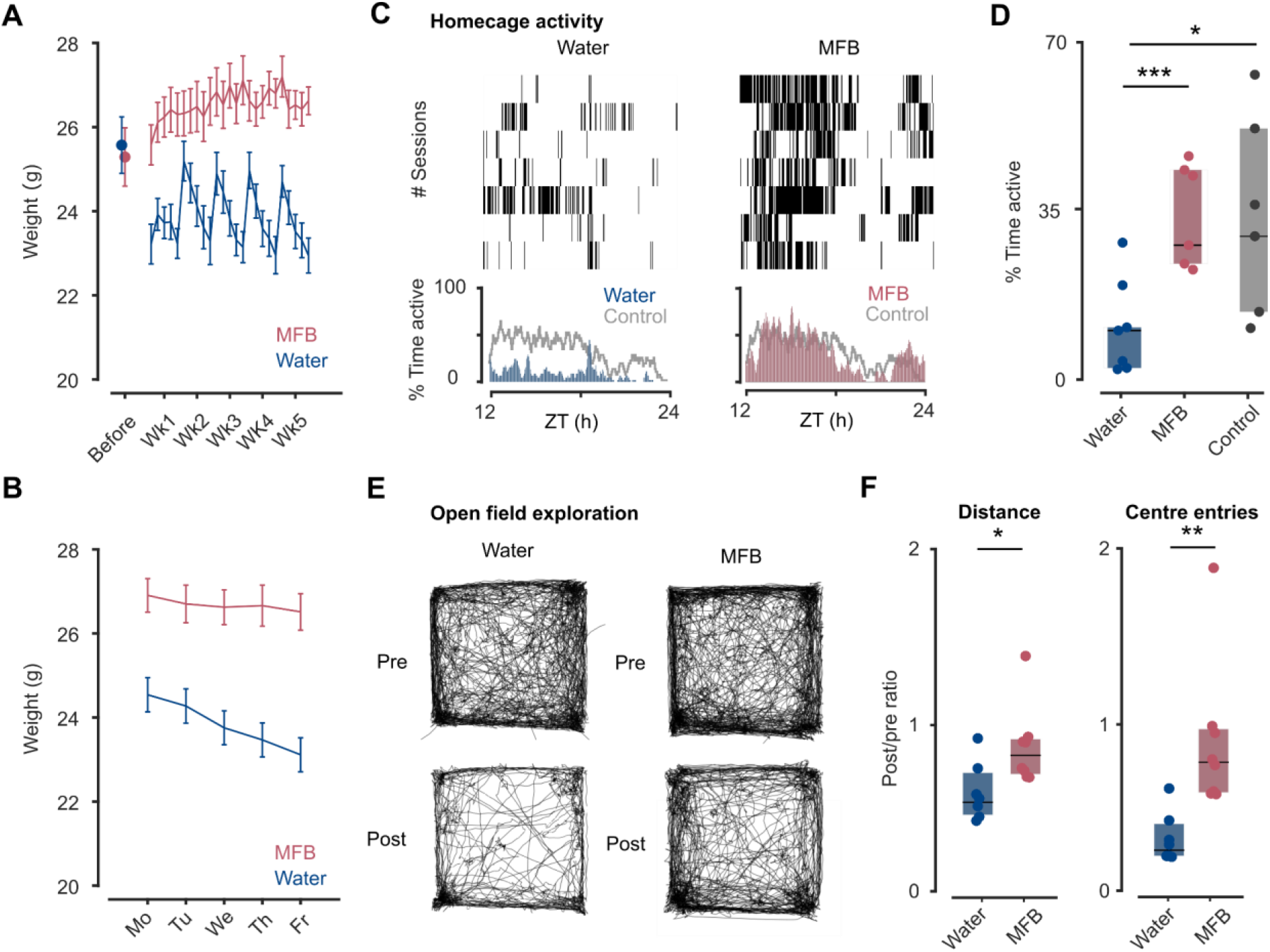
Improved mouse well-being using MFB stimulation relative to water deprivation protocol. A. Weight evolution starting from the week prior to any behavioural experiments (isolated point) and across the first five weeks during which mice were trained five days a week. (n=7, 8 mice, error bar are sem). In this panel and following, only data from MFB lever pressing mice was collected. B. Weight evolution averaged across days of the week (n=7, 8 mice, error bar are sem). C. Top : temporal distribution of periods of activity during the lights-off period for each session for a cage containing two animals (black bar indicates a period during which wheel running was detected). Bottom : mean activity throughout the lights-off period for MFB and water-deprived mice (blue and red) with the mean activity of control mice (gray) with no interventions overlayed D. Percent of time spent active throughout the whole night for MFB and water deprived animals as well as control animals with no interventions (Wilcoxon rank sum test, pval : 0.007, 0.02 ranksum statistic : 73, 58, n=7,7,6 sessions) E. Superposed trajectories for mice during the open field session. F. Ratio of total distance travelled (left) and number of entries into the centre of the field (right) before and after training during 5min exploration of the open field (Wilcoxon ranksum test, pval : 0.039, 0.028, ranksum statistic : 83,89, n=7,8 mice)

Overall, we have shown that the use of the MFB stimulation does not induce any motor agitation and or any increase in impulsive behaviour relative to the water deprivation approach. Moreover, it attenuates the impact of learning protocols on markers of stress relative to water deprivation. The novel protocol is therefore compatible with the experimental demands of well-controlled and calm behaviour and can contribute to increased animal well-being.

### Response latency and discrimination times in MFB stimulated mice

We finally assessed whether the MFB reward task reproduces subtler aspects of behaviour such as response times. In a recent report, it was shown that discrimination time is longer for the difficult PTvsFM task than for the easier PTvsPT task [8]. We applied the same methodology to our data to estimate the earliest time at which the Go and the No-go sound could be significantly distinguished based on the animal’s response (Fig. 6A illustrates the method). Consistent with previous observations established using water-deprivation, we found longer discrimination times for the PTvsFM task for both MFB-reward and water deprivation (Fig. 6 B,D). Moreover there was a good quantitative match between the two methods despite using differences in reward and motor response (median for MFB and water: PTvsPT: 240/220ms, PTvsFM: 490/580ms).

**Figure 6.**
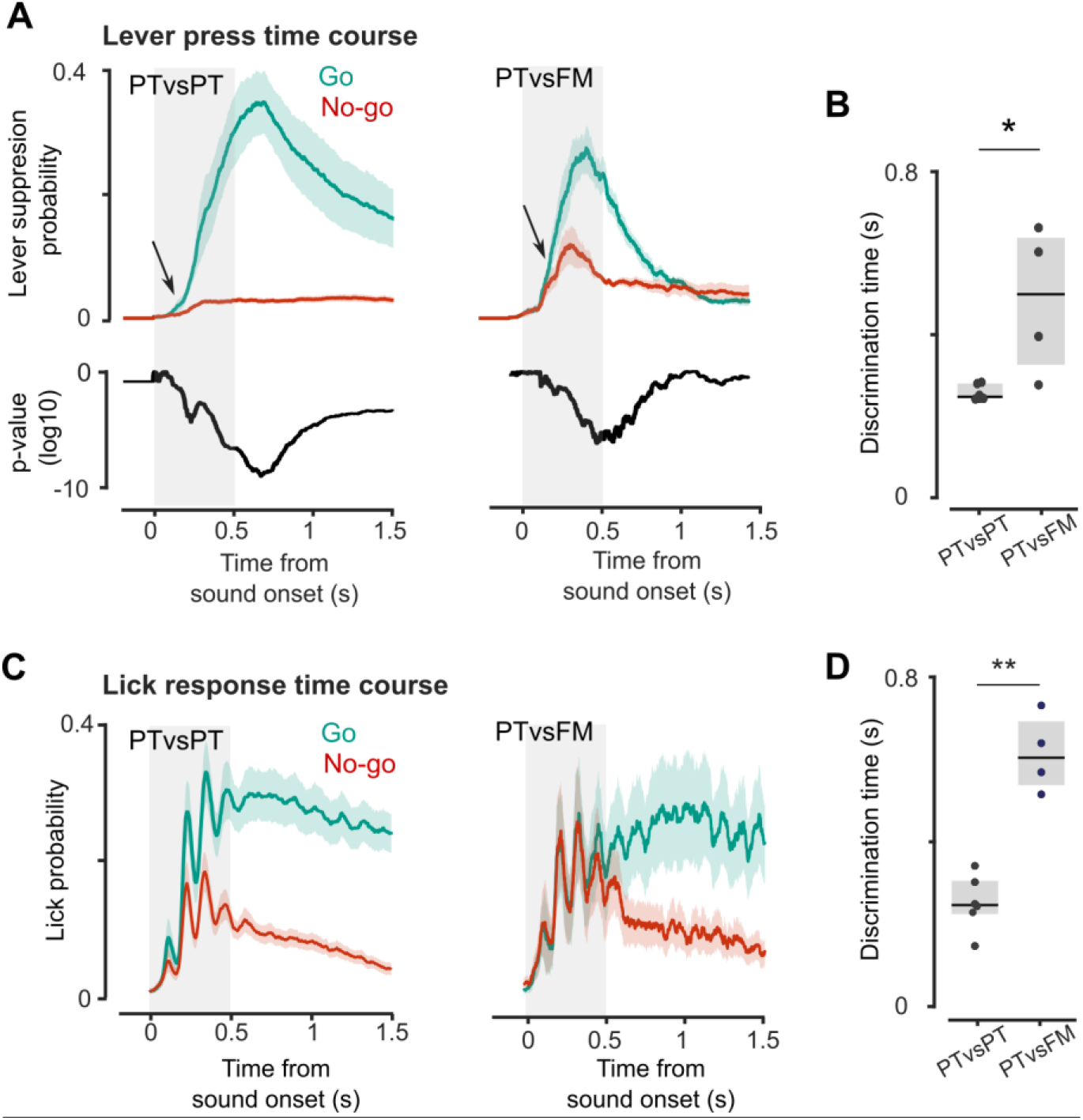
Similar task dependance of discrimination times for MFB and water-reward protocols. A. Average lever press probability for PTvsPT and PTvsFM tasks across all go and No-go trials. Arrow shows the estimated discrimination time defined as the earliest time at which p-value of a t-test for the difference between the go and No-go traces (bottom pannel) is below 0.01. Error bars are sem. B. Discrimination time as defined in A for each animal. (n= 8,4 Wilcoxon rank sum test, pval = 0.03, ranksum statistic : 23) C. Average lick probability for PTvsPT and PTvsFM tasks across all go and No-go trials. D. Discrimination time, defined as earliest time where Go and No-go lick signals are significantly different, for the PTvsPT and PTvsFM tasks (n=7, 4, Wilcoxon rank sum test, pval = 0.001, ranksum statistic : 23, only animals that reached critertion accuracy for PTvsFM task are included)

We also found in MFB-rewarded mice that response times were significantly shorter for false alarm No-go trials than for Go trials (Fig. S6A,B) in accordance with the literature [17]–[19]. We reasoned that the difference was driven by early, non-discriminative responses perhaps associated with an excessive motivation to perform reward collection. If this is the case, we would expect behavioural epochs with short lever press latencies on Go trials to be associated with low discrimination accuracy (i.e. high FA rates). Indeed, we found that FA rate and lever press latencies in Go trials were positively correlated (Fig. S6C-E).

These results exemplify how our MFB paradigm can be easily transferred to the study of perceptual discrimination since it reproduces known results from the literature including subtler aspects of behaviour such as temporal dynamics. Therefore, our direct stimulation of the reward system appears to leave intact much of the core mechanisms underlying perception and decision-making, allowing for their continued study with this new approach.

## Discussion

By implementing stimulation of the MFB as a reward in mice, we have shown in various auditory behavioural tasks that this approach provides an efficient and high-yield protocol for learning without changing psychometric measures (Fig. 2G) or response dynamics (Fig. 6). MFB stimulation allows to train animals for thousands trials per session without loss of motivation, which can reduce the number of training days, especially for more difficult tasks (Fig. 2). MFB stimulation removes the need to adjust deprivation levels to control motivation. This more easily yields a fast and reliable learning outcome across animals and high steady state performance level as we observed when comparing our MFB-stimulation and water deprivation protocols in a difficult task (Fig. 2). MFB rewarded mice also showed reduced impulsivity and signs of stress compared to their water deprived counterparts, which have previously been reported as having modified behaviour [20], [21].

The strongest advantage of MFB stimulation is the lack of satiation, providing high and stable motivation throughout long sessions. Increased trial counts per session is an asset for psychometric measurements to increase the number of stimulus variations tested, and also for protocols involving the study of neural activity. Indeed, it allows to collect data from the same neurons whilst providing the freedom to examine neural responses to variations in stimuli or task contingency. It is therefore important to stress that MFB stimulation is compatible with recordings of neural data as shown in [4] and [5]. Moreover, we observed no stimulation-induced agitation which is important when combined with imaging techniques that require field-of-view stability.

MFB stimulation provides a novel methodology to train mice in conditions that are conducive to their well-being, when compatible with the underlying scientific question. In particular, the approach refines current protocols by minimising the impact of the experiment on mice’s daily activities since MFB mice are not deprived in any way. The drawback of the MFB protocol is that it requires the implantation of an extra electrode during head-post surgery. In behavioural experiments that do not require surgeries, the potential stress of this additional procedure may counterbalance the benefits.

There are limitations in the comparisons we made between MFB stimulation and water-deprived protocols. Although the two tasks were operated by the same experimenters we cannot ensure that all aspects of the protocols were similarly optimised. In MFB-rewarded mice, we have tested two motor reports (lever pressing and licking) and only one for water deprived mice. Lever pressing requires motor learning [7] contrary to licking. This requirement slowed learning of sound-response association phase (Go training) for lever pressing compared to licking, and may have also an impact on learning rates in the discrimination task. Another difference between the protocols is that the water-deprived animals discriminated Go from No-go stimuli by exceeding a threshold on the number of licks whereas MFB-rewarded mice were constrained to an all or none response (1 lever press / 1 lick). This is also likely to impact learning speed. The criterion used for water deprived mice being more permissive with respect to impulsivity, it is likely that water deprived mice were given an easier protocol than MFB mice. We therefore believe that the conclusion that MFB stimulation provides a more stable setting enabling faster acquisition at the cohort level in difficult tasks remains valid in practice, despite implementation discrepancies.

A potential issue with direct stimulation of the reward system is that it is a less ethologically relevant motivation than water or food and therefore could distort both behavioural and neural activity. It is important to bear in mind that states of deprivation, although ethologically relevant, are not “neutral” states. They induce metabolic changes [20], most obvious in weight loss, behavioural stress, as we measured (Fig. 5.E-F), and brain-wide changes in neural activity [21]. A recent study showed that food-deprivation induces energy saving changes in neural coding which lead to a slight impairment of visual discrimination both at the neuronal and behavioural performance level [22]. These findings are likely also relevant for water-restriction since it leads to reduced food-intake. In this study, we have shown that psychometric performance (Fig. 2) and response timing (Fig. 6, Fig. S6) are very similar between water and MFB reward. Therefore, for the majority of studies focused on sensory or motor behaviours and not motivation or reward per se, MFB stimulation is unlikely to modify the results of interest and provides a way to avoid the side-effects of deprived states.

MFB stimulation evokes dopamine release in the Nucleus Accumbens (NA) [23] and pharmacologically increasing or reducing dopamine levels increases or reduces MFB stimulation efficacy respectively [24]. Although the MFB contains dopaminergic fibres essential for instrumental behaviour [25], the physiology of MFB self-stimulation has been shown to be incompatible with a direct recruitment of these fibres to drive reward [26]. Instead, the rewarding level of stimulation is best predicted by the level of direct activation of another set of currently undefined, descending, myelinated fibres in the MFB. It is hypothesised that these fibres indirectly drive dopamine release [26].

Fine comparisons of optogenetic dopaminergic release with MFB stimulation highlight differences between the two paradigms [27] and large forebrain lesions including the NA leave MFB self-stimulation intact [25]. The MFB also houses glutamatergic, monoaminergic and GABAergic fibres that extend to around 50 different brain areas that are both antidromically and orthodromically activated upon stimulation [26]. Therefore, a complex array of neural systems beyond the NA participate in the effect of MFB stimulation and they remain to be dissected [28], [29].

MFB stimulation is clearly distinguished from other rewards by the absence of satiety and the remarkable stability of its rewarding valence. Factors that lead to satiation for other rewards (ex sucrose accumulation in the gut for food reward) do not impact MFB stimulation efficacy [30]. The MFB reward can lead to seemingly ‘pathological’ behaviours with large stimulation intensity, during which animals prefer to self-stimulate for MFB reward over all other choices, leading to maladaptive behaviours such as self-starvation (preference of MFB stimulation over food) [31], [32][33]. Importantly, these behaviours only occur with ad libitum access to strong MFB stimulation. We therefore did not witness any such behaviours in MFB-rewarded mice trained in our lab. Calibration of voltage (Fig1 D.G) to use the minimal efficient voltage is likely beneficial to reduce such negative consequences. In line with this, parametric studies which evaluate the impact of different factors on response-intensity curves similar to those used in our calibration protocol show that MFB reward in fact can compete with food reward and the rewarding effect of both MFB and food can summate [34]. These results suggest that the MFB stimulation and more ethological rewards are evaluated along shared dimensions. The commonalities between MFB stimulation and other rewards is supported by neuronal recordings. First, during self-stimulation the evoked dopamine release is reduced when stimulation is expected [35], [36]. Therefore dopamine release is maximal when reward is unexpected in accordance with reward prediction error as found for other rewards [37]. Second, neurons in the hypothalamus respond with similar changes in firing pattern to MFB stimulation and to glucose [38]. Third, changes in brain reward circuitry such as chronic food-restriction or leptin administration impact both MFB stimulation and food intake [39].

However MFB stimulation has a wider range of effects than ethological rewards. Stimulation that is not contingent on reward invigorates responding [40] and post-training simulation can enhance memory [41]. Although these extra effects may positively impact learning in sensory stimulation tasks, experimenters should bear in mind that transitioning from water or food reward to MFB stimulation is therefore not just a change in reward paradigm but may have broader impacts.

A number of other studies have proposed refinements of the water deprivation protocol. Adding sucrose to water can increase trial number [42] and replacing the animal’s drinking water with ad libitum access to citric acid water allows for a less impactful but still efficient version of the deprivation protocol [11], [43]. However, even with these improvements mice still undergo weight loss due to deprivation and deprivation methods so far do not allow to obtain more than a thousand trials per session (Fig. 3D, Table 1). Recently, Burgess et al. [9] demonstrated that optogenetic activation of dopaminergic neurons in the ventral tegmental area could be used as a very efficient reward system. This approach showed similar advantages to ours in terms of accelerating learning and enhancing trial yield. However, electrical MFB stimulation is more flexible and easier to implement. First, optogenetic stimulation requires the use of a specific CRE line making it incompatible with studies in WT or other CRE lines of interest to the researcher. Second, whereas MFB implantation is achieved with one brief surgery, the optogenetic approach requires injecting a viral vector and waiting for expression (2-3 weeks) before finally implanting the optic fibre in a second surgery.

Here we explored only a few specific tasks amongst the many available. Further work would be required to expand the technique to different sensory modalities and task rules. It is likely that MFB will further generalise given that it has already been shown to work well in freely moving animals both with nose-poke [5] and with spatial conditioning [4], [6]. Finally, further optimisation of MFB stimulation parameters (delay, duration, etc) could lead to improved results.

In conclusion, MFB stimulation adds to the toolbox of methods available to train rodents depending on the demands of the scientific question. It is an easy-to-implement alternative to deprivation and the strong improvement in trial-yield that it provides should enable and accelerate novel experiments that require learning fine perceptual measurements, or complex task rules, or that aim at linking single-trial behaviour to neural data.

## Materials and methods

### Subjects and surgical procedures

7 to 9-week old C57Bl6 male mice (Janvier Labs) were used. Mice were housed in an animal facility (08:00–20:00 light) in cages of 3 to 5 individuals with access to enrichment (running wheel, cotton bedding and wooden logs).

All experiments were performed in accordance with the French Ethical Committee (Direction Générale de la Recherche et de l’Innovation) and European legislation (2010/63/EU). Procedures were approved by the French Ministry of Education in Research after consultation with the ethical committee #59 (authorization number APAFIS#28757-2020122111048873).

Surgeries were performed in order to equip all mice with a metal head-plate for fixation. Implantations of a bipolar stimulation electrode (60-μm-diameter twisted stainless steel, PlasticsOne) in the medial forebrain bundle were performed for the MFB group. Mice were injected with buprenorphine (Vétergesic, 0,05-0,1 mg/kg) 30min prior to anaesthesia induction with isoflurane (3%) and then placed on a thermal blanket during the whole procedure with isoflurane maintained at 1-1.5%. The eyes were protected with Ocrygel (TVM Lab). Lidocaine was injected under the skin of the skull 5 minutes prior to incision.

The MFB electrode was implanted using stereotaxic coordinates (AP -1.4, ML +1.2, DV +4.8) (Fig. S1 for histological verification). The electrode was then fixed along with the headplate to the skull using dental cement (Ortho-Jet, Lang).

After surgery, mice received a subcutaneous injection of glucose (G30) and metacam (1mg/kg). Mice were subsequently housed for one week with metacam delivered via drinking water or dietgel (ClearH20) without any manipulation. All animals were co-housed before and after surgery without any degradation to the implants.

6 mice were housed in identical conditions to provide a control group for night time activity recordings. These mice were left in their cages during the whole period prior to activity recordings which were performed at the same age as for traine mice.

### Behavioural setup

Behavioural experiments were monitored and controlled using homemade software (Elphy, G. Sadoc, UNIC, France for water deprivation and Matlab homemade GUI for MFB) coupled to a National Instruments card (PCIe-6351). Sounds were amplified (SA1 Stereo power amp, Tucker-Davis Technologies) and delivered through high frequency loudspeakers (MF1-S, Tucker-Davis Technologies) in a pseudo-random sequence. Mice were placed on a 3D printed contention tube with two piezo-electric sensors inserted at the base to allow monitoring of movement.

#### Water deprivation

Water delivery (5μl/reward) was controlled with a solenoid valve (LVM10R1-6B-1-Q, SMC). A voltage of 5V was applied through an electric circuit joining the lick tube and aluminium foil on which the mouse was sitting, so that lick events could be monitored by measuring the voltage through a series resistor in this circuit.

#### MFB stimulation

MFB stimulation was delivered via a pulse train generator (PulsePal V2, Sanworks) that produced 2ms biphasic pulses at 50Hz for 100ms. *Lever group*: The stimulation was delivered when the mouse pressed the lever, raising the opposite end and interrupting an infrared beam (ZHITING, infrared motor speed metre, 3.3V). The lever was weighted to fall back into its initial position as soon as the mouse removed its paw. It was designed using Autodesk Fusion 360 and produced with an Ultimaker-3 3D printer. *Licking group* : MFB-rewarded mice were using the same setup as water-deprived mice.

### Behavioural procedures

Fig. S3 provides an overview of the behavioural protocols.

#### Sounds and task structure

Mice were trained on three different tasks. In each case, sounds of 500ms were presented at 70dB with a sample rate of 192kHz. The identity of Go and No-go sounds was counterbalanced across mice.

#### Frequency discrimination task (PTvsPT)

Discrimination between a pure tone at high frequency (16kHz) and a pure tone at low frequency (6kHz).

#### Frequency psychometrics (PTvsPT PS)

Discrimination between a group of 8 high frequency pure tones (10kHz -16kHz) and a group of 8 low frequency pure tones (6kHz - 9.5kHz) in which the 16 tones were equally spaced on a logarithmic scale.

#### Pure tone vs frequency-modulated sound (PTvsFM)

Discrimination between a pure tone at 6kHz and either a frequency sweep rising from 6 to 16kHz or a 2-step rise consisting of 250ms of 6kHz and 250ms of 16kHz.

#### MFB stimulation

Mice performed behaviour five days per week (Monday to Friday). During the entire behavioural training period, food and water were available *ad libitum*. Animal weight was monitored daily.

#### Habituation

During four days, mice were habituated to head fixation by progressively increasing the duration from 30min to 2 hours without reward or sound stimulation. Free reward

#### (Lever press)

During 1 or 2 days mice received systematic reward when they pressed the lever with a 2s minimal inter reward interval. In order to encourage lever pressing, the experimenter actively shaped the mouse’s behaviour for 10-50 trials by placing its paw on the lever with a small plastic stick. During this initial phase, before calibration, stimulation started at 3V and was reduced by the experimenter if it elicited a motor reaction.

#### Free reward (licking)

During a short 30 min session, mice received a systematic reward when they licked the spout with a 500ms inter reward interval. In order to encourage licking as the mice are not water-deprived, the spout is placed very close to the mouse’s mouth. During this initial phase, before calibration, stimulation started at 3V and was reduced by the experimenter if it elicited a motor reaction.

#### Go training

Once animals were spontaneously pressing or licking, go training was conducted. Go trials were presented with 80% probability, while the remaining trials were blank trials (no stimulus). A trial consisted of a random inter-trial interval (ITI) between 1 to 2 s to avoid prediction of stimulus appearance, a random ‘no press’/’no lick’ period between 3 and 5 s and a fixed response window of 1.5 s. For the lever group, any lever press longer than 50ms was counted as a lever press. For the lick group, a single lick was registered as a response. Lever press/lick during the response window on a Go trial was scored as a ‘hit’ and triggered an immediate MFB stimulation. No lever press/lick was scored as a ‘miss’ and the next trial immediately followed.

#### Calibration

Once reliable report to the sound was established (>80% hits for the Go stimulus), the voltage of the stimulation was systematically reduced from the voltage used during go training down to 0 in 0.5V increments in order to obtain a calibration curve. During all following experiments, the minimal voltage that elicited >90% of maximum pressing/lickins was used. We found that this voltage could be adjusted by 0.5V to either motivate (increase) the mouse during harder tasks or to slow unsolicited response (decrease) and that this could facilitate learning. Calibration was performed using the Go sound from the PTvsPT protocol since this was learnt first. For mice using licking report, this calibration can also be performed before Go sound training by simply measuring the number of spontaneous licks at different voltage levels.

#### Go/No-go training

After Go training, the second sound (No-go) was introduced. During presentation of the No-go sound, the absence of lever press/lick was scored as a ‘correct rejection’ (CR) and the next trial immediately followed. Any report during No-go trials was scored as a ‘false alarm’ (FA), no stimulation was given, and the animal was punished with a random time-out period between 5 and 7 s. Each session contained 50% probability for each trial type.

#### Psychometric measurement

Once mice reached 80% accuracy on the PTvsPT task, the number of sounds increased to 16 (8 high frequency sounds and 8 low frequency sounds) in order to establish psychometric curves for frequency discrimination (PTvsPT - PS task).

During all phases of the protocol, the session duration was 1h30-3h and was only limited by time constraints on the use of the apparatus.

#### Criteria defining mice with functional MFB stimulation (licking)

Given that mice rapidly and spontaneously licked to receive MFB stimulation we evaluated implantation success during the “Free reward” phase (Fig. S2B). Mice were counted as positively responding if within a maximum of two 30min sessions they reached licking levels >0.25licks/s.

#### Criteria defining mice with functional MFB stimulation (lever pressing)

Given that lever pressing is a complex motor behaviour that takes a few days to master and that conditioning can be confused with a simple increase in movement during which the mouse randomly activates the lever, we evaluated implantation success during the “Go training” phase (Fig. S2F). Mice were counted as positively responding if they reached >80% accuracy in this training phase within a maximum of 10 days. Mice that showed no progress during more than 5 days were counted as non-functional without performing the full 10 days of training.

#### Impact of inter-trial interval

We measured whether MFB rewarded mice could maintain their performance at high levels even with a short inter-trial interval. We performed in succession on the same day 4 bouts of 200 Go/No-go trials, decreasing the average total inter-trial interval (random ITI + random no press window) from an average of 5s down to 0.5s.

#### Maximal number of trials measurement

To provide a benchmark for the number of trials obtainable in a single session we performed on the PTvsPT task a 4 hour session with a 2s ITI. Mice were checked at regular intervals during the session for signs of stress and offered water after 2h.

#### Coton interaction assay

Before beginning training on the task, the efficacy of MFB stimulation can be rapidly tested using an object interaction assay in order to eliminate non-responding mice. We performed stimulation at 3V. We performed this in freely moving mice in a neutral cage with a cotton swab. After an initial 5 min habituation phase, mice undergo 5 min conditioning during which when they touch the swab they receive stimulation (Fig. SG). During conditioning the preference is generally obvious by eye (Video 1). To quantify the impact of the stimulation we manually timed the time spent interacting with the swab during the habituation and stimulation phase (Fig. S2H-I). Mice which fail to increase their interaction time with the swab can be tested at a higher voltage (we have tried up to 6V with no adverse reactions) and if no clear preference is manifest then the stimulation is counted as non-efficient.

#### Water deprivation

Mice performed behaviour five days per week (Monday to Friday). Water deprivation began after habituation to head fixation. Before starting the training procedure, mice were water restricted for two consecutive days. Afterwards, water restriction (0.8mL/day) was interleaved with a 16h *ad libitum* supply overnight every Friday. During the entire behavioural training period, food was available ad libitum. Animal weight was monitored daily.

#### Habituation

Prior to deprivation, during four days, mice were habituated to head fixation by progressively increasing the duration from 30min to 2 hours without reward or sound stimulation.

#### Free lick

During the first day of training, mice received water by licking the lick port without any sound.

#### Go training

After this period, Go training was conducted where Go trials were presented with 80% probability, while the remaining trials were blank trials (no stimulus). A trial consisted of a random inter-trial interval (ITI) between 3 to 0 s to avoid prediction of stimulus appearance, a random ‘no lick’ period between 1.5 and 3.5 s and a fixed response window of 1.5s. Licking during the response window on a Go trial above lick threshold (3-5 consecutive licks) was scored as a ‘hit’ and triggered immediate water delivery. Licking below the threshold was scored as a ‘miss’ and the next trial immediately followed. At the beginning of each session, 10 trials with ‘free rewards’ were given independently of the licking to motivate the mouse.

#### Go/No-go training

When animals reached more than 80% ‘hits’ for the Go stimulus, the second sound (No-go) was introduced, and the lick count threshold was set between 5 and 8 licks (this parameter was adjusted at the beginning of training for each mouse). During presentation of the No-go sound, licking below threshold was scored as a ‘correct rejection’ (CR) and the next trial immediately followed, licking above threshold on No-go trials was scored as a ‘false alarm’ (FA), no water reward was given, and the animal was punished with a random time out period between 5 and 7 s. Each session contained 40% probability for the Go and No-go stimuli and 20% blank trials.

#### Psychometric measurement

Once mice reached 80% accuracy on the PTvsPT task, the number of sounds increased to 16 (8 high frequency sounds and 8 low frequency sounds) in order to establish psychometric curves for frequency discrimination (PTvsPT - PS task).

During all phases of the protocol, the session was continued until animals had received a maximum of 0.8mL or until they ceased licking in response to sound. This occurs within 300-600 trials. Throughout the protocol, we performed 320 trials per day except towards the end of training on Thursday or Friday when mice sometimes performed up to 640 trials. Mice that did not attain 0.8mL during the session received the missing amount directly in their cage

#### Open field

Before the start of learning and then again after 3 months of training, mice were placed at the centre of a brightly lit 50cmx50xm open field for three minutes. The surface and walls were cleaned between each mouse. The animal’s position was tracked using an online homemade tracking program (Matlab, Mathworks).

#### Home cage activity

In order to measure mice’s activity level in their homecage, we monitored the turning of a running wheel from 8 pm to 8 am, the dark period which was undisturbed by behavioural experiments. The measurement was made by counting infrared beam breaks due to the passage of an opaque section of the wheel. We counted as “active” any period of 5s during which the wheel made one full turn.

#### Histology

In order to extract the brain for histology, mice were deeply anaesthetised using a ketamine-medetomidine mixture. Once all reflexes had disappeared we performed an electrolytic lesion using a 20s of continuous 5V stimulation. Mice were then perfused intracardially with 4% buffered paraformaldehyde fixative. The brains were carefully dissected and left in paraformaldehyde overnight and then sliced into 100um sections using a vibratome.

### Data analysis

#### Learning and performance analysis

Learning curves were obtained by calculating the fraction of correct responses over blocks of 200 trials. Accuracy over one session was calculated as (hits + correct rejections)/total trials. The learning rate is quantified by the number of training days to reach accuracy greater than or equal to 80%. Note that for all evaluations of learning time we only took into account valid trials left after excluding all periods during which the mouse failed to respond for more than 15 trials.

#### Response time analysis

Response times are defined as the time to first lever press / first lick. Discrimination time was defined as the earliest 10ms time bin at which a significant difference between lever pressing / licking signal for Go and No-go trials was found. We used the non-parametric Wilcoxon rank-sum test to obtain the p-value for the null hypothesis that the Go and No-go distributions were identical for each time bin and then applied the Benjamini-Hochberg correction for multiple testing over all time bins. In order to avoid biassing statistical significance due to differing number of trials, we performed multiple random subsampling of datasets to restrict to 100 trials and took the mean reaction time over all shuffles.

#### Piezoelectric signal

The acquired piezoelectric signal was filtered to remove 50Hz contamination. Changes in the animal’s position within the tube can create baseline shifts in the signal. We therefore measured the amount of movement using the absolute value of the first-order temporal derivative. The absolute values of this signal cannot be compared between MFB-rewarded and water-deprived mice since water-deprived mice were placed on aluminium foil in order to detect licking and the deformations of the foil attenuate the values readout by the sensor.

#### Statistics

For each statistical analysis provided in the manuscript, the Kolmogorov–Smirnov normality test was first performed on the data. If the data failed to meet the normality criterion, statistics relied on non-parametric tests. We therefore represent the median and quartiles of data in boxplots in all figures, in accordance with the use of non-parametric tests. Ranksum and signed rank: we report the signed rank statistic if the number of replicates is too weak to provide the normal Z statistic

### Code and 3D printer files availability

In order to facilitate other laboratories trying out the new methodology, we have made available on a Github page (https://github.com/AntoninVerdier/MFBasic) the 3D printer files for contention tube and lever, simplified Arduino code to implement a minimalist Go/No-go task with MFB stimulation and a detailed description of all the hardware needed and how to set it up. For further assistance, please contact the corresponding author.

## Acknowledgements

We thank Léa Vinel for assistance with behavioural experiments.

## Author contributions

SB contributed the original idea of the project and designed research with AV and BB. SB and AV performed experiments and analysed the data. Mouse training was co-supervised by SB and AV and performed by ND and DG. AA designed the lever-pressing detection and helped with experiments. SB and AV analysed the data and wrote the paper with comments from BB. BB provided funding and equipment for task implementation.

## Supplementary materials

**Figure S1.**
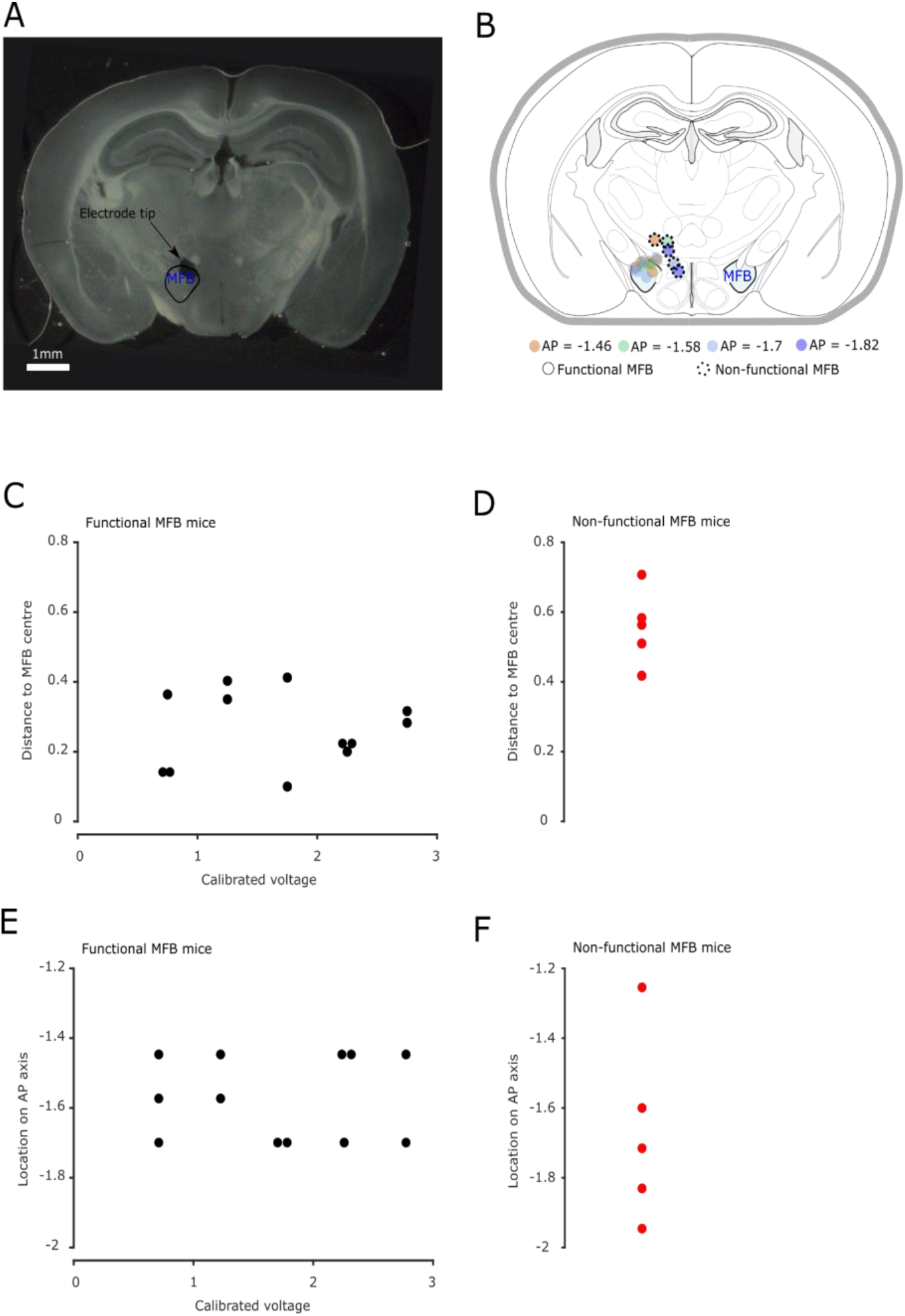
MFB stimulation electrode localisation. A. Post-hoc localisation of electrode tip via electrolytic lesion in an example mouse. B. Localisation of electrode tips from multiple mice. Note that implantation sites for mice without functional MFB (as defined based on learning of report behaviour - see methods and FigS2) are well outside the region. Note that not all animals were involved in the auditory discrimination tasks studied in this manuscript in order to have a sufficient number of mice with non-functional MFB. See table S1 for breakdown of mice in each experiment. (n=17) C. Distance of electrode tip to the centre of the MFB as a function of the voltage used after calibration. No correlation was observed (Pearson correlation coefficient : R=0.03, pval = 0.91, n=17). D. Distance of electrode tip to the centre of the MFB without functional MFB. E.F. As in C and D but for location along the A/P axis. (Pearson correlation coefficient : R=0.0 pval = 1, n=17).

**Figure S2.**
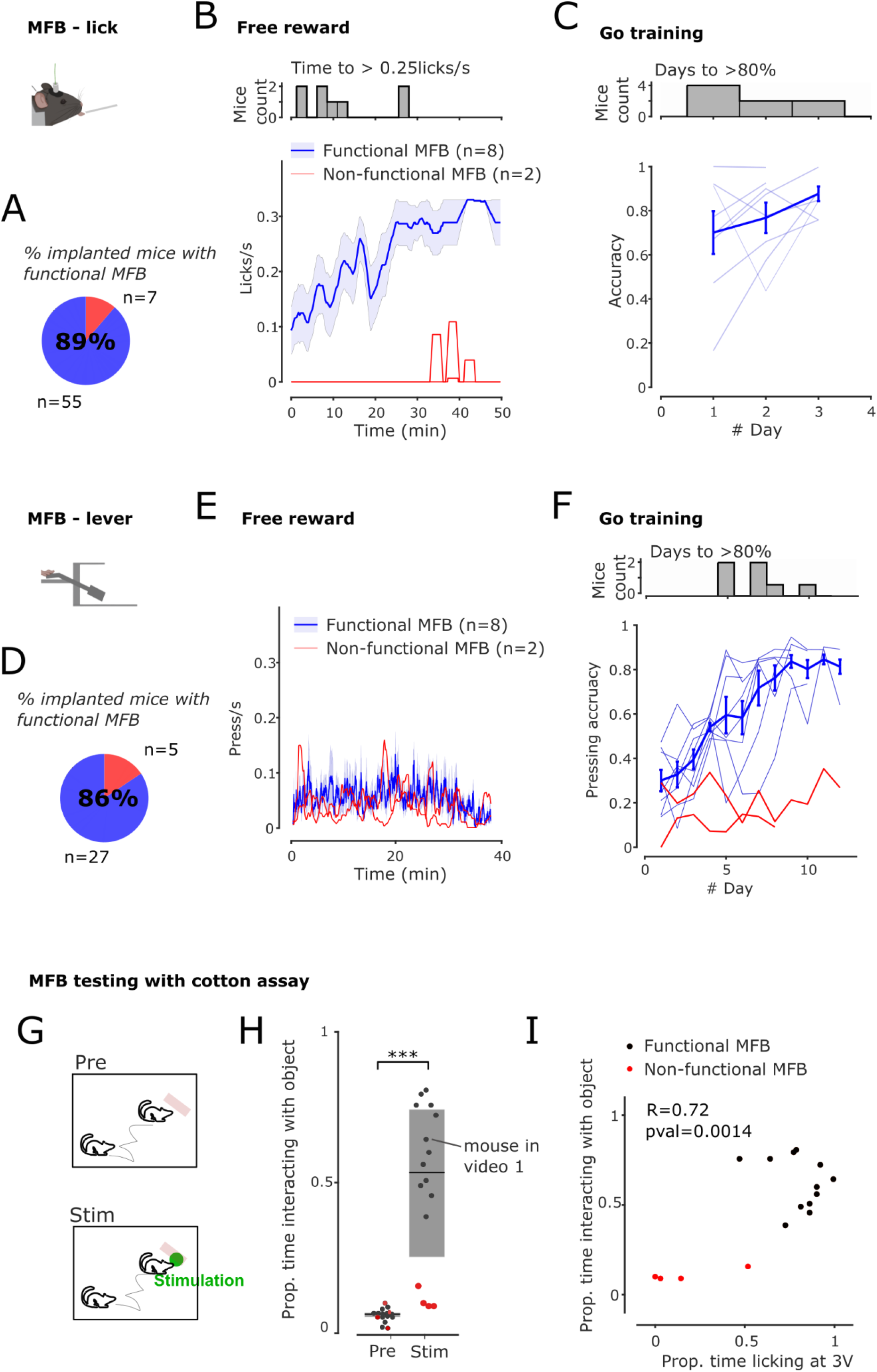
Acquisition of motor response and MFB efficacy testing. A. Proportion of functional MFB implantations for all mice performing a licking task implanted in the laboratory (defined according to criterion in methods and illustrated by blue and red lines in S2B). Note that not all animals were involved in the auditory discrimination tasks studied in this manuscript but include all mice implanted in our laboratory to provide a robust statistic. See table S1 for breakdown of mice in each experiment. All mice with functional MFB stimulation went on to successfully learn the auditory tasks on which they were trained. B. Licking rates during the initial free-lick phase for mice with functional (blue, n=8) and non-functional (red, n=2) MFB stimulation. C. Daily accuracy in the Go training phase of the PTvsPT task showing that mice reliably learnt to lick to the Go sound within 1-2 days. D. Proportion of functional MFB implantations for all mice performing a lever pressing task implanted in the laboratory (defined according to criterion in methods and illustrated by blue and red lines in S2E-F). E. Lever press rates during the initial free press phase of mice with functional (blue, n=8) and non-functional (red, n=2) MFB stimulation. It should be noted that this phase does not allow to clearly discriminate between the two groups of mice, this may be due to the overall low pressing rates and the fact that in mice with poorly situated MFB electrodes, stimulation can increase motor activity leading to non-specific activation of the lever. F. Daily accuracy in the Go training phase of the PTvsPT task showing that mice required around 7 days of training to develop a reliable lever suppression strategy in response to the sound. During this phase, mice with non-functional MFB stimulation can be clearly identified by the absence of any increase in accuracy.. G. Schematic of cotton assay. Interaction time with a cotton pellet is measured during 5minutes without and then with 3V MFB stimulation every time the mouse was in contact with the cotton swab. H. Interaction time with the cotton swab strongly increased in all mice, except those with non-functional MFB stimulation. (Wilcoxon signed rank test, pval = 4.3E-4, zval: -3.51, n = 16) I. Proportion of time interacting with cotton swab versus proportion of time licking at the same voltage during calibration showing that the cotton assay has good predictive power of the efficacy of MFB stimulation as a reward to drive licking behaviour. (Pearson correlation : R=0.72, pval = 0.0014, n=16)

**Figure S3.**
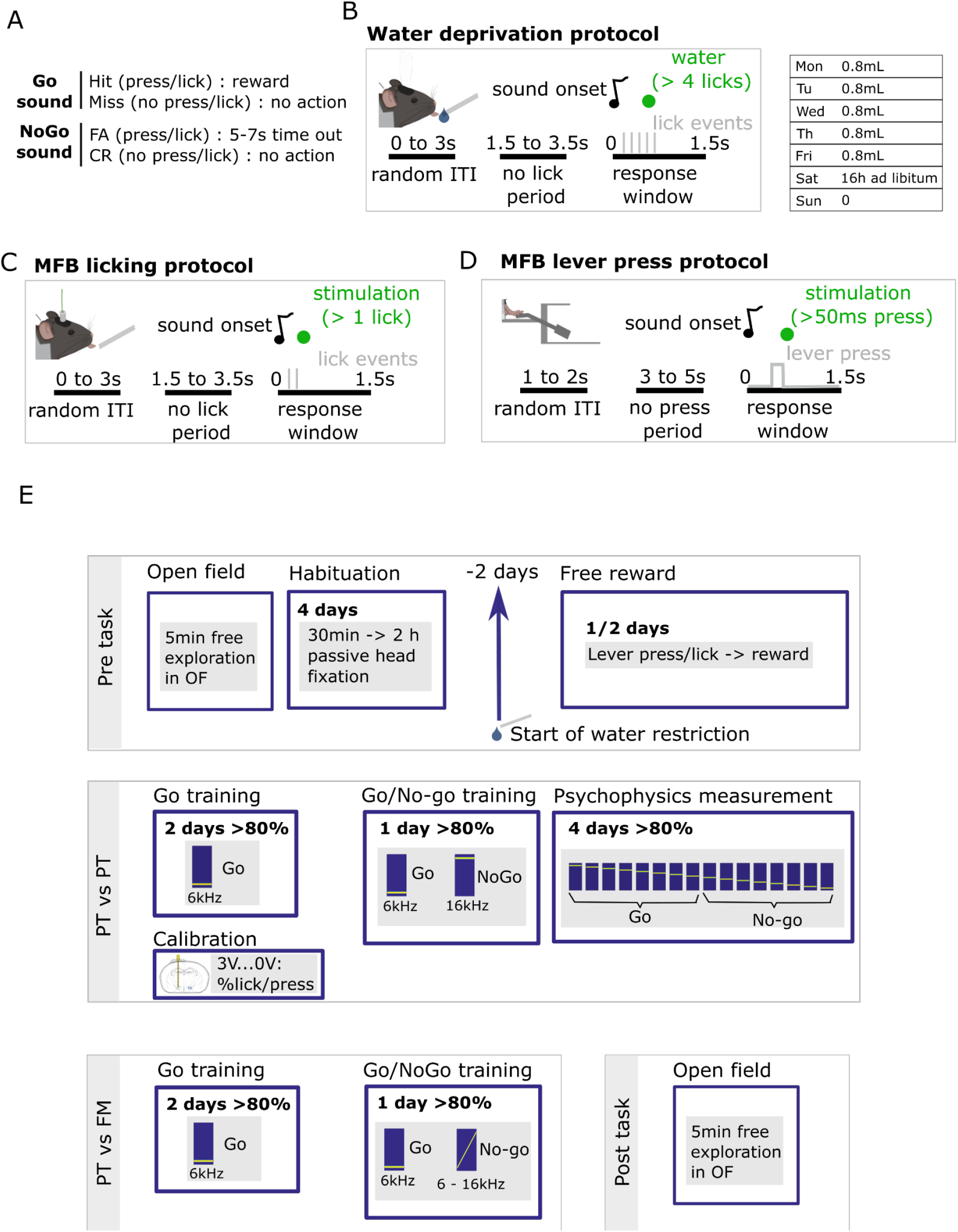
Behavioural protocols. A. Definition of trial types for all reward and response contingencies. B. Timing of events in discrimination task for water deprived animals and weekly water restriction regimen. Here we use an example threshold of 4 licks but this was fixed at the beginning of training depending on the animal’s behaviour. C. Timing of events in discrimination task for MFB rewarded animals that used licking as a report. D. Timing of events in discrimination task for MFB rewarded animals that used lever pressing as a report. E. Steps of the behavioural protocol outlining the succession of tasks. All information applies to both MFB-rewarded and water-deprived animals unless indicated by the corresponding pictogram. Note that open field tests were only performed on lever pressing MFB mice.

**Figure S4.**
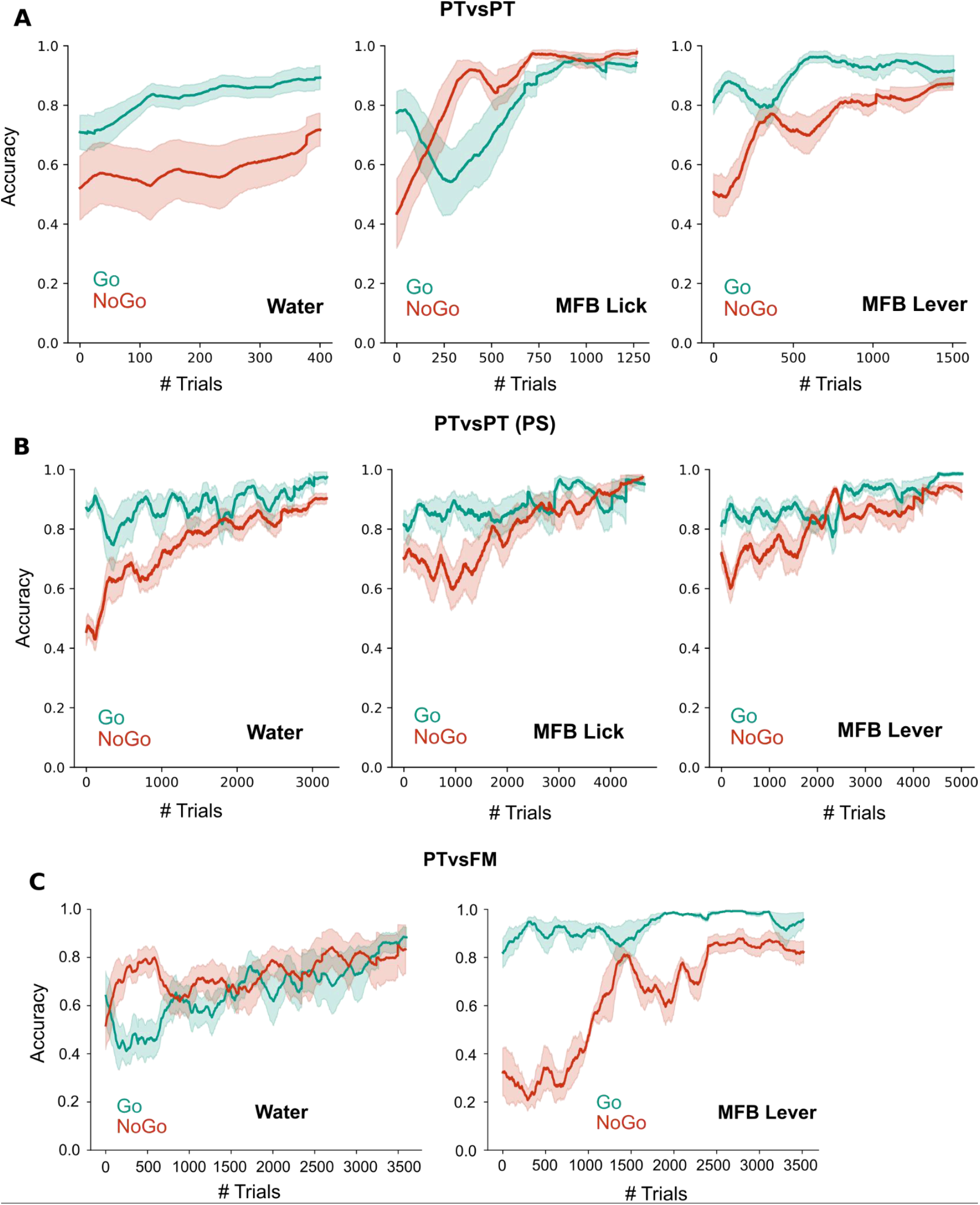
Evolution of Go / No-go accuracy during training in all tasks. A. B. C. Learning curves for Go and No-go trials for water-deprived, MFB lick and MFB lever groups performing the pure tone (PTvsPT) discrimination task (A - n = 7,8,8), the pure tone psychometric task (PTvsPT - PS) requiring classification of high and low pure tones (B - n = 7,8,8) and the pure tone vs frequency modulated (PTvsFM) discrimination task (C - n = 7,4). MFB lever PTvsPT (5.75 +/- 1.6 sessions, 375 +/- 91 trials per session) MFB lever PTvsPT - PS (9 +/- 2.5 sessions, 831+/-224 trials per session) MFB lever PTvsFM (9.7 +/- 4.5 sessions, 552+/-55 trials per session) MFB lick PTvsPT (4.5 +/- 2 sessions, 536 +/-272 trials per session) MFB lick PTvsPT - PS (6.8 +/- 1.5 sessions, 1035 +/-101 trials per session) Water deprived PTvsPT (4.75 +/- 1.6 sessions, 320 +/-0 trials per session) Water deprived PTvsPT - PS (16.7 +/- 6.3 sessions, 300 +/-22 trials per session) Water deprived PTvsFM (21 +/- 8 sessions, 390 +/-50 trials per session)

**Figure S5.**
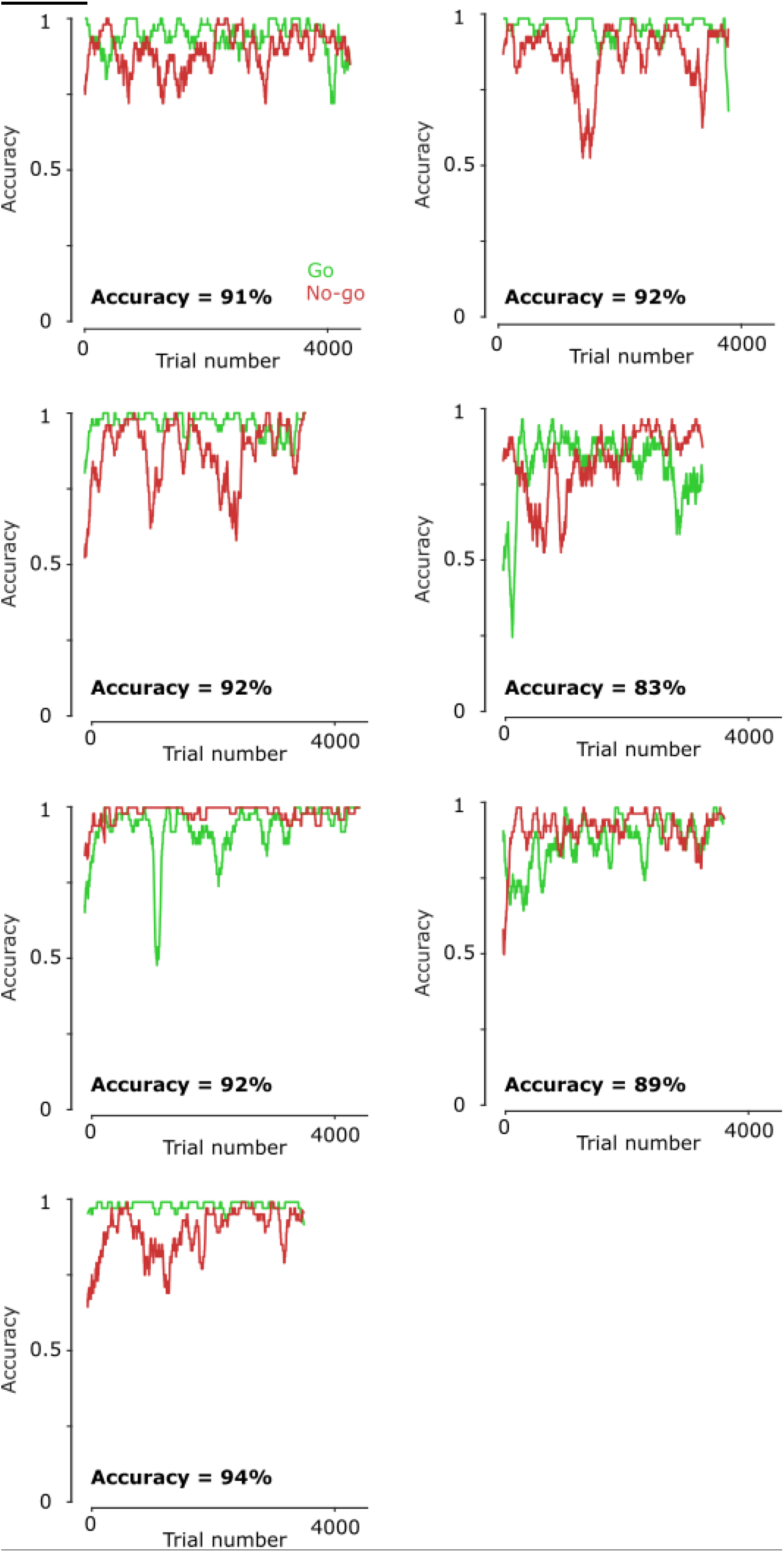
Full sessions for all mice during 4h sessions with fast ITI. Each pannel shows the go and No-go accuracy for one mouse during a 4H session performing the PTvsPT task.

**Figure S6:**
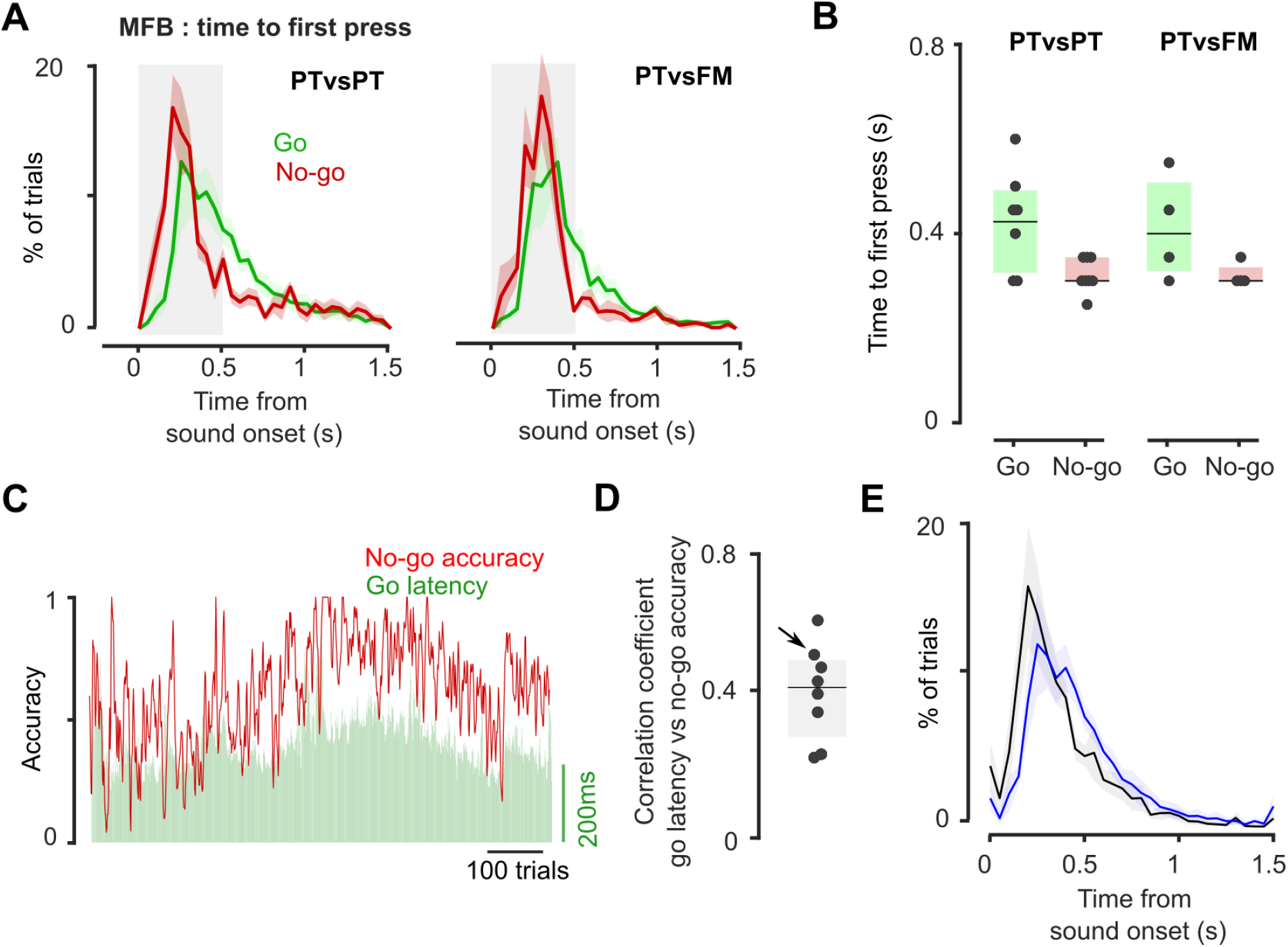
Link between reaction time and task accuracy. A Histogram of reaction times (defined as time to first lever press) for PTvsPT and PTvsFM tasks on correct go trials and incorrect No-go trials (MFB : n=7 & 4, error bars are SEM). B. Reaction times for MFB groups (one mouse out of eight for PTvsPT was excluded as an outlier using the criteria of three scaled MAD from the median) C. Concatenated blocks of trials for an example mouse during which go accuracy was consistently over 95% showing the correlation between fluctuations in No-go accuracy (red) and the go latency (green). D. Correlation coefficient between No-go accuracy (red) and the go latency (green) for all mice. (n=8 mice, pearson correlation coefficient, all p-values<1E-4) Arrow indicates the animal shown in C. E. Histogram of reaction times to Go stimulus for blocks with low No-go accuracy (<60%, black) and high No-go accuracy (>70%, blue) showing that, during blocks in which animals reliably pressed for No-go trials, lever presses on Go trials occurred later.

**Table S1:** Overview of mice used in the study

|  |  |
| --- | --- |
| Water deprivation (PTvsPT, PTvsPT-PS & PTvsFM) | n=7 for PTvsPT, PTvsPT-PS & PTvsFM |
| MFB lever press (PTvsPT, PTvsPT-PS & PTvsFM) | n=8 for PTvsPT PTvsPT-PS & subgroup n=4 for PTvsFM |
| MFB licking (PTvsPT, PTvsPT-PS) | n=8 for PTvsPT PTvsPT-PS |
| Histology (Fig. S1) | n = 17 (8 from task used in paper + 9 from other tasks in lab) |
| Coton test / calibration comparison (Fig. S2G) | n = 16 (8 from task used in paper + 8 from other tasks in lab) |
| Total MFB lever press for success rate estimate (FigS2A) | n = 32 (8 from task used in paper + 24 from other tasks in the lab)<br>27/32 responding : 86% |
| Total MFB licking for success rate estimate (FigS2D) | n = 62 (8 from task used in paper + 54 from other tasks in lab)<br>55/62 responding : 89% |

**Video 1 : Example mouse in cotton assay**

**Video 2 : Example mouse performing the Go training phase of PTvsPT by lever pressing**

**Video 3 : Example mouse performing the Go-No-go training phase of PTvsPT task by licking**

